# High-resolution species delimitation in *Acinetobacter baumannii* using a novel Core-Gene Consensus Delimitation approach

**DOI:** 10.64898/2026.05.01.722318

**Authors:** Khaoula El Mchachti, Adam Valcek, Charles Van der Henst, Jean-François Flot

**Affiliations:** Evolutionary Biology & Ecology, Department of Organismal Biology, Université libre de Bruxelles (ULB), Brussels, Belgium; Interuniversity Institute of Bioinformatics in Brussels – (IB)2, Brussels, Belgium; Microbial Resistance and Drug Discovery, VIB-VUB Center for Structural Biology, VIB, Flanders Institute for Biotechnology, Brussels, Belgium; Structural Biology Brussels, Vrije Universiteit Brussel (VUB), Brussels, Belgium

**Author notes:** Address correspondence to Khaoula El Mchachti, Charles Van der Henst, and Jean-François Flot,.

## Abstract

*Acinetobacter baumannii* is a highly adaptable nosocomial pathogen with extensive antibiotic resistance, a disproportionately large accessory genome, and high genomic plasticity. Owing to these features, the World Health Organisation (WHO) classifies *A.baumannii* as a critical-priority pathogen. In this study, we analyzed 47 isolates from our VUB (Vrije Universiteit Brussel) collection and applied distance-based species-delimitation algorithms – Automatic Barcode Gap Discovery (ABGD) and Assemble Species by Automatic Partitioning (ASAP) – for the first time at the bacterial core-genome scale. By integrating conspecificity matrices, we extended these traditionally single-locus methods into a multi-locus framework, which we term Core-Gene Consensus Delimitation (CGCD).

Across a range of gene-level co-occurrence thresholds, CGCD consistently recovered 11 stable groups using both ABGD and ASAP. Larger-scale validation using 856 *A. baumannii* genomes recovered the same 11 well-separated groups were recovered, demonstrating the robustness and reliability of our clustering approach. Mapping these groups onto a core-genome phylogeny revealed that each group forms a distinct clade, indicating that they represent evolutionarily independent lineages rather than arbitrary clusters. We further constructed a clustering tree based on accessory gene presence–absence patterns. In this tree, only one strain (AB231-VUB) clustered within group 11; otherwise, the groups remained tightly cohesive, sharing characteristic sets of accessory genes.

Together, these results show that the groups defined by CGCD are genomically, evolutionarily, and functionally distinct, supporting their interpretation as separate species. Our findings highlight CGCD as a powerful, high-resolution framework for species delimitation. CGCD is threshold-free, gene-based, and universally applicable—the first species-delimitation approach that can be applied across all domains of life, from bacteria to animals and plants.

## INTRODUCTION

*Acinetobacter baumannii* is one of the most alarming nosocomial pathogens worldwide, frequently causing severe infections in intensive care units and showing high levels of antibiotic resistance [1, 2]. Clinically, *A. baumannii* infections are associated with high morbidity and mortality rates, owing to its extensive drug resistance, robust biofilm formation, and ability to persist on hospital surfaces under adverse conditions. These features make it a major cause of ventilator-associated pneumonias, bloodstream infections, and other serious clinical complications [3, 4]. Compounding the clinical challenge, *A. baumannii* has remarkable genetic plasticity (driven by a large accessory genome and pervasive horizontal gene transfer [5, 6]), which complicates its taxonomic delineation.

Historically, species delimitation in many fields of biology has relied heavily on single-locus analyses. From a single genetic marker, researchers have delineated species using three main approaches: tree-based, distance-based and allele sharing-based approaches [7, 8]. Among these, the distance-based approaches, such as Automatic Barcode Gap Discovery (ABGD) and its upgrade Assemble Species by Automatic Partitioning (ASAP) identify discontinuities (gaps) in distance distributions, without the need for setting an a priori delimitation threshold [9, 10]. These methods are widely popular in studies of eukaryotes such as animals and fungi as they are very fast and easy to understand, but are less used in prokaryotic taxonomy. They are quite conservative and tend to overlump rather than oversplit taxa, i.e. they fail to distinguish highly similar species rather than erroneously consider conspecific populations or strains as two different species [11].

In bacteria and archaea, single-locus analyses have long been dominated by the 16S rRNA gene, valued for its highly conserved regions and universal amplification across prokaryotes [12]. However, 16S rRNA gene alone can be misleading in defining species boundaries for *A. baumannii* [13, 14, 15]. In particular, many bacterial genomes harbor multiple 16S rRNA gene copies, sometimes divergent enough to appear as separate species if examined in isolation [13]. Furthermore, the slow evolutionary rate of 16S rDNA often fails to capture the subtle genetic differences crucial for delineating closely related strains [15]. Thus, while 16S rDNA remains a helpful baseline marker, it underestimates the genomic plasticity observed in *A. baumannii*, which can exhibit substantial variation outside its conserved core genome [6, 16].

To overcome these limitations, the field has moved toward multilocus and whole-genome approaches. In studies of eukaryotes, multilocus methods based on the multispecies coalescent model, such as BPP (which stands for Bayesian Phylogenetic and Phylogeography [17]), are very slow when handling more than a handful of sequences and tend to confound intraspecific population structure with species boundaries [18]. An alternative is to run single-locus species delimitation methods on several loci and integrated the result into a consensus delimitation using a conspecificity matrix [19]. In prokaryotes, similarly, multi-locus sequence typing (MLST) provided a more nuanced view of strain relationships than 16S only and has facilitated global epidemiological tracking [20, 21]. More recently, whole-genome similarity measures such as Average Nucleotide Identity (ANI) have revolutionized bacterial systematics [22, 23]. Genomes sharing an ANI above 95–96% are generally considered conspecific, with stricter thresholds (up to 98–99.5%) now being explored to detect subspecies-level variation and cryptic population structures [24, 25]. However, ANI remains constrained by fixed similarity thresholds and its reliance on average nucleotide identity across shared genomic regions, which can obscure gene-level variation and differences in accessory genome content. ANI estimates are also sensitive to genome quality, are computationally demanding in large-scale comparisons, and are limited in their ability to resolve recent divergence, recombination dynamics, and ecologically meaningful population structure. As a result, closely related strains with distinct evolutionary or functional traits may remain indistinguishable under ANI-based frameworks.

Inspired by eukaryotic species delimitation methods, we propose a different strategy that leverages the availability of large numbers of complete genome for bacteria and archaea: instead of limiting ABGD or ASAP to a single marker or to a handful of them, we apply them both to each gene in the *A. baumannii* core genome. Each gene produces an independent set of strain groupings. From these, we build a conspecificity matrix recording, for every pair of strains, how many times they are placed together across all core genes. By scanning different co-occurrence thresholds, we identify stable and biologically coherent clusters – effectively turning ABGD and ASAP into a genome-wide consensus “voting” system. We then compare the resulting delimitation with those obtained from running ABGD and ASAP on a single concatenated core-genome alignment, as well as with a clustering tree based on presence/absence of individual acccessory genes.

This Core-gene Consensus Delimitation (CGCD) framework is, to our knowledge, the first systematic, core-genome-scale application of ABGD and ASAP in bacteria. By combining the conservative, gap-detection logic of these methods with genome-wide resolution, CGCD offers a robust, reproducible means to resolve fine-scale population structure in a clinically important pathogen.

## MATERIALS AND METHODS

### Bacterial strains

The 47 *A. baumannii* genomes used in this study were downloaded in March 2024 from the Acinetobase collection [26] hosted at VUB, which compiles well-characterized clinical isolates exhibiting multidrug-resistant (MDR), extensively drug-resistant (XDR), pandrug-resistant (PDR), or carbapenem-resistant phenotypes. These 47 VUB strains were recovered through a national microbiological surveillance program conducted in Belgian hospitals and were subsequently whole-genome sequenced using both Illumina MiSeq (short-read) and Oxford Nanopore (long-read) platforms (Table **1**).

**TABLE 1.** Characteristics of datasets of *A. baumannii* used in our analyses.

| Strain | Nanopore SRA | Nanopore data (Gb) | Average Nanopore Q score | Illumina SRA | Illumina data (Gb) | Average Illumina Q score |
| --- | --- | --- | --- | --- | --- | --- |
| AB5075-VUB | SRR13700038 | 1.6 | 29.19 | SRR13700039 | 0.1787 | 30.00 |
| AB169-VUB | SRR17842164 | 0.8066 | 12.63 | SRR18189635 | 0.3878 | 20.47 |
| AB227-VUB | SRR17842135 | 0.9842 | 13.47 | SRR14718264 | 0.4838 | 20.17 |
| AB229-VUB | SRR17842134 | 1.1 | 13.47 | SRR14718286 | 0.2954 | 20.53 |
| AB232-VUB | SRR17842131 | 1.1 | 13.47 | SRR14718266 | 0.3145 | 19.98 |
| AB177-VUB | SRR17842158 | 1 | 12.55 | SRR14718275 | 0.2465 | 20.71 |
| AB186-VUB | SRR17842152 | 0.6554 | 16.24 | SRR14718277 | 0.357 | 20.42 |
| AB21-VUB | SRR17842126 | 1.3 | 13.68 | SRR18189636 | 0.2353 | 20.19 |
| AB40-VUB | SRR17842122 | 0.6721 | 12.62 | SRR14718288 | 0.4264 | 19.89 |
| AB179-VUB | SRR17842157 | 1.4 | 12.60 | SRR18189634 | 0.3834 | 19.92 |
| AB32-VUB | SRR17842125 | 1.4 | 13.70 | SRR14718268 | 0.283 | 20.05 |
| DSM30011-VUB | SRR17842127 | 1.4 | 12.29 | SRR18189637 | 0.2637 | 20.80 |
| ATCC17978-VUB | SRR17842129 | 1.4 | 12.31 | SRR18189638 | 0.2233 | 20.08 |
| ATCC19606-VUB | SRR17842128 | 1.2 | 12.20 | SRR18189639 | 0.2278 | 20.13 |
| AB176-VUB | SRR17842159 | 1.1 | 12.57 | SRR14718274 | 0.2369 | 20.07 |
| AB188-VUB | SRR17842150 | 0.6421 | 16.13 | SRR14718280 | 0.3731 | 20.11 |
| AB20-VUB | SRR17842133 | 1.6 | 13.72 | SRR14718287 | 0.2784 | 19.77 |
| AB187-VUB | SRR17842151 | 0.5233 | 16.23 | SRR14718279 | 0.404 | 19.62 |
| AB14-VUB | SRR17842155 | 1.3 | 13.69 | SRR14718278 | 0.2228 | 19.99 |
| AB9-VUB | SRR17842166 | 0.5437 | 13.70 | SRR14718271 | 0.2018 | 19.00 |
| AB231-VUB | SRR17842132 | 0.805 | 13.46 | SRR14718265 | 0.2684 | 20.67 |
| AB181-VUB | SRR17842154 | 0.5602 | 16.22 | SRR14718257 | 0.269 | 20.73 |
| AB3-VUB | SRR17842167 | 0.8081 | 13.68 | SRR14718252 | 0.2529 | 20.01 |
| AB171-VUB | SRR17842163 | 0.8204 | 12.62 | SRR14718272 | 0.3301 | 20.55 |
| AB172-VUB | SRR17842162 | 1 | 12.54 | SRR14718273 | 0.3001 | 20.64 |
| AB36-VUB | SRR17842124 | 0.7748 | 12.60 | SRR14718269 | 0.2853 | 20.20 |
| AB175-VUB | SRR17842160 | 1.2 | 12.60 | SRR14718256 | 0.269 | 20.42 |
| AB212-VUB | SRR17842146 | 0.6708 | 16.11 | SRR14718282 | 0.4198 | 18.63 |
| AB193-VUB | SRR17842148 | 0.5259 | 16.32 | SRR14718251 | 0.319 | 20.10 |
| AB213-VUB | SRR17842145 | 0.6362 | 16.16 | SRR14718283 | 0.3831 | 19.18 |
| AB173-VUB | SRR17842161 | 1.3 | 12.60 | SRR14718255 | 0.2808 | 20.57 |
| AB226-VUB | SRR17842136 | 1.2 | 13.42 | SRR14718263 | 0.2759 | 20.06 |
| AB183-VUB | SRR17842153 | 0.4778 | 16.20 | SRR14718258 | 0.3048 | 20.65 |
| AB180-VUB | SRR17842156 | 0.6943 | 16.27 | SRR14718276 | 0.3109 | 20.21 |
| AB214-VUB | SRR17842143 | 0.5415 | 16.22 | SRR14718259 | 0.5289 | 18.11 |
| AB233-VUB | SRR17842130 | 1.1 | 13.42 | SRR14718267 | 0.2425 | 20.58 |
| AB217-VUB | SRR17842141 | 1.5 | 13.47 | SRR14718284 | 0.4346 | 18.50 |
| AB224-VUB | SRR17842137 | 1.1 | 13.46 | SRR14718262 | 0.386 | 21.47 |
| AB194-VUB | SRR17842147 | 0.6083 | 16.23 | SRR14718281 | 0.4084 | 19.78 |
| AB16-VUB | SRR17842144 | 1.4 | 13.65 | SRR14718253 | 0.2866 | 20.54 |
| AB220-VUB | SRR17842139 | 1.1 | 13.45 | SRR14718285 | 0.4366 | 19.00 |
| AB216-VUB | SRR17842142 | 1.3 | 13.43 | SRR14718260 | 0.4714 | 18.40 |
| AB219-VUB | SRR17842140 | 1 | 13.43 | SRR14718254 | 0.3928 | 17.81 |
| AB189-VUB | SRR17842149 | 0.5318 | 16.42 | SRR14718250 | 0.3807 | 19.49 |
| AB222-VUB | SRR17842138 | 1.1 | 13.42 | SRR14718261 | 0.4846 | 19.34 |
| AB167-VUB | SRR17842165 | 1.1 | 12.62 | SRR14718289 | 0.2848 | 20.77 |
| AB39-VUB | SRR17842123 | 1.1 | 12.62 | SRR14718270 | 0.2388 | 19.52 |

The assembly of the genomes of each strain was described previously [27]; the characteristics of the resulting assemblies are summarized in Table **2**.

**TABLE 2.** Characteristics of the *A. baumannii* assemblies used in our analyses.

| Strain | Assembly accession | Assembly size (Mb) | Nb of contigs | N50 (Mb) | Completeness | Contamination |
| --- | --- | --- | --- | --- | --- | --- |
| AB5075-VUB | GCA_016919505.2 | 4.1 | 4 | 4 | 100 | 0.08 |
| AB169-VUB | GCA_022459415.1 | 4.1 | 1 | 4.1 | 100 | 0.19 |
| AB227-VUB | GCA_022467655.1 | 3.9 | 1 | 3.9 | 100 | 0.08 |
| AB229-VUB | GCA_022467535.1 | 3.9 | 1 | 3.9 | 100 | 0.09 |
| AB232-VUB | GCA_018788735.1 | 3.8 | 587 | 0.0156 | 100 | 0.07 |
| AB177-VUB | GCA_022459095.1 | 4 | 1 | 4 | 100 | 0.22 |
| AB186-VUB | GCA_018789495.1 | 3.9 | 316 | 0.046 | 100 | 0.18 |
| AB21-VUB | GCA_022460295.1 | 4.1 | 1 | 4.1 | 100 | 0.24 |
| AB40-VUB | GCA_018789005.1 | 3.9 | 276 | 0.0321 | 100 | 0.28 |
| AB179-VUB | GCA_022459075.1 | 3.6 | 1 | 3.6 | 100 | 0.08 |
| AB32-VUB | GCA_018788855.1 | 3.9 | 359 | 0.0248 | 100 | 0.08 |
| DSM30011-VUB | GCA_022466815.1 | 4 | 1 | 4 | 100 | 0.02 |
| ATCC17978-VUB | GCA_022467135.1 | 3.9 | 1 | 3.9 | 100 | 0.16 |
| ATCC19606-VUB | GCA_022466975.1 | 4 | 1 | 4 | 100 | 0.06 |
| AB176-VUB | GCA_018789355.1 | 4.5 | 1,064 | 0.013 | 100 | 0.13 |
| AB188-VUB | GCA_018789475.1 | 3.9 | 402 | 0.0225 | 100 | 0.16 |
| AB20-VUB | GCA_018789235.1 | 4 | 513 | 0.0154 | 100 | 0.11 |
| AB187-VUB | GCA_018789615.1 | 3.9 | 337 | 0.0298 | 100 | 0.16 |
| AB14-VUB | GCA_018789595.1 | 3.9 | 394 | 0.0227 | 100 | 0.18 |
| AB9-VUB | GCA_018791845.1 | 3.8 | 479 | 0.0158 | 100 | 0.12 |
| AB231-VUB | GCA_022467435.1 | 4 | 1 | 4 | 100 | 0.13 |
| AB181-VUB | GCA_022548575.1 | 4.1 | 1 | 4.1 | 100 | 0.3 |
| AB3-VUB | GCA_018788905.1 | 3.7 | 405 | 0.0187 | 100 | 0.03 |
| AB171-VUB | GCA_022459295.1 | 4 | 1 | 4 | 100 | 0.12 |
| AB172-VUB | GCA_022459175.1 | 4 | 1 | 4 | 100 | 0.1 |
| AB36-VUB | GCA_022460015.1 | 4 | 1 | 4 | 100 | 0.12 |
| AB175-VUB | GCA_022459135.1 | 3.9 | 1 | 3.9 | 100 | 0.12 |
| AB212-VUB | GCA_022468565.1 | 3.9 | 1 | 3.9 | 100 | 0.1 |
| AB193-VUB | GCA_022468755.1 | 3.9 | 1 | 3.9 | 100 | 0.05 |
| AB213-VUB | GCA_022468455.1 | 3.9 | 1 | 3.9 | 100 | 0.11 |
| AB173-VUB | GCA_022459155.1 | 4.1 | 1 | 4.1 | 100 | 0.07 |
| AB226-VUB | GCA_022467735.1 | 4 | 1 | 4 | 100 | 0.04 |
| AB183-VUB | GCA_022469075.1 | 4 | 1 | 4 | 99.99 | 0.23 |
| AB180-VUB | GCA_022459055.1 | 3.9 | 1 | 3.9 | 100 | 0.05 |
| AB214-VUB | GCA_022468355.1 | 4 | 1 | 4 | 100 | 0.14 |
| AB233-VUB | GCA_022467255.1 | 3.9 | 1 | 3.9 | 100 | 0.07 |
| AB217-VUB | GCA_022468195.1 | 4.1 | 1 | 4.1 | 100 | 0.21 |
| AB224-VUB | GCA_022467805.1 | 4 | 1 | 4 | 100 | 0.17 |
| AB194-VUB | GCA_018789335.1 | 3.9 | 261 | 0.0314 | 100 | 0.12 |
| AB16-VUB | GCA_022460575.1 | 4 | 1 | 4 | 100 | 0.11 |
| AB220-VUB | GCA_022467975.1 | 4 | 1 | 4 | 100 | 0.2 |
| AB216-VUB | GCA_022468295.1 | 3.9 | 1 | 3.9 | 100 | 0.09 |
| AB219-VUB | GCA_022468075.1 | 3.9 | 1 | 3.9 | 100 | 0.2 |
| AB189-VUB | GCA_022468795.1 | 4 | 1 | 4 | 100 | 0.12 |
| AB222-VUB | GCA_022467905.1 | 3.9 | 1 | 3.9 | 100 | 0.07 |
| AB167-VUB | GCA_022459595.1 | 4 | 1 | 4 | 100 | 0.09 |
| AB39-VUB | GCA_022459875.1 | 4 | 1 | 4 | 100 | 0.13 |

### Genomic analyses

The assembled genomes were first annotated using Prokka (v1.14.6) [28], which generated GFF annotation files for each isolate. These annotation files were subsequently processed with Roary (v3.12.0) [29] using default parameters to construct a gene presence/absence matrix. This matrix categorized genes into core genes (shared by all isolates) and accessory genes, which may provide isolates with adaptive advantages such as antibiotic resistance or enhanced virulence.

### Applying species delimitation methods

To determine species boundaries within our VUB *A. baumannii* dataset, we employed two computational methods: Automatic Barcode Gap Discovery (ABGD) and Assemble Species by Automatic Partitioning (ASAP). ABGD detects “barcode gaps” within distance distributions to automatically partition sequences into putative species clusters, while ASAP expands on this by using a distance-based scoring system to select the best species partitions [9, 10].

### 16S rRNA alignment and concatenated core-genome alignment

We applied ABGD and ASAP to two different datasets. The first dataset consisted of 16S rRNA alignments, representing a traditional single-locus marker commonly used in bacterial taxonomy. We extracted and retained all copies of the 16S rRNA gene annotated in each genome to capture intra-genomic variability. The second dataset consisted of a single concatenated alignment of all core genes shared across isolates.

### Average nucleotide identity (ANI)

We used FastANI (v1.31) to calculate the mean nucleotide identity between the genomes of the analyzed isolates [30]. Pairwise ANI results were parsed and formatted into a symmetrical matrix using a custom Python script. Hierarchical clustering was performed on a distance matrix derived from ANI values (100 - ANI), and the clustered matrix was visualized as a heatmap using the seaborn library.

### Core-gene consensus delimitation (CGCD)

Only genes present in all isolates were retained to ensure a consistent dataset. Each of these core genes was aligned independently using MAFFT v7 [31] under default parameters.

Following alignment, the two distance-based species delimitation algorithms –ABGD and ASAP– were applied separately to each core gene. Each algorithm generated gene-specific partitions, showing how isolates were grouped together at different loci. For each core gene, we retained the most frequently observed ABGD partition and the partition with the best (smallest) ASAP score. From these results, we created a binary partition matrix for each core gene, where rows and columns corresponded to strains, and each cell contained a value of 1 if the pair of strains was placed in the same group for that gene, and 0 otherwise. These matrices captured the grouping pattern for every gene individually.

We then combined all per-gene partition matrices by summing them to produce a single conspecificity matrix [19].To define final clusters, we applied a co-occurrence threshold to the conspecificity matrix. We scanned thresholds corresponding to different proportions of the total number of core genes, starting at 50% and increasing to 100%. For each threshold, strains were grouped if they co-clustered in at least that proportion of core genes, and the resulting number of groups was recorded. This process was visualized in a curve plotting the number of groups against the applied threshold. The curve exhibited a stability plateau, indicating a range of thresholds over which group composition remained unchanged. From this plateau, we retained the stable number of groups as the final delimitation outcome.

### Large-scale genome dataset

Following the identification of genomic groups based on the dataset of 47 VUB *A. baumannii* strains, a large-scale analysis was conducted to assess the effect of expanded genome sampling on our genomic delimitation framework. A dataset comprising 856 publicly available *A. baumannii* genomes was retrieved from the NCBI database, including all genomes available at the time of download. These genomes were processed using the same CGCD framework. This large-scale analysis was used to evaluate the scalability of our CGCD approach when applied to a substantially larger genomic dataset.

### Phylogenetic and accessory genes analysis

The core genome generated by Roary was used to to infer a phylogenetic tree using FastTree v2.1.11 [32]. In parallel, the accessory gene clustering tree produced by Roary was retained for comparative analysis. Both the core-genome phylogeny and the accessory gene clustering tree were visualized using the online tool iTOL (Interactive Tree Of Life) [33], and groups identified from previous species delimitation analysis were mapped onto both trees to compare clustering patterns between core and accessory genome signals.

## RESULTS

### Pan-genome analysis

The presence/absence matrix generated by Roary shows the distribution of genes across all isolates and provides an overview of gene conservation and variability within the pangenome(Figure **1**). We categorized the genes into four groups: core genes (present in 46–47 isolates), soft-core genes (present in 44–46 isolates), shell genes (present in 7–44 isolates), and cloud genes (present in fewer than 7 isolates) (Figure **2**). The core genes made up only 25% of the total genome, while 75% were classified as accessory genes (soft-core, shell, or cloud).

**FIG 1.**
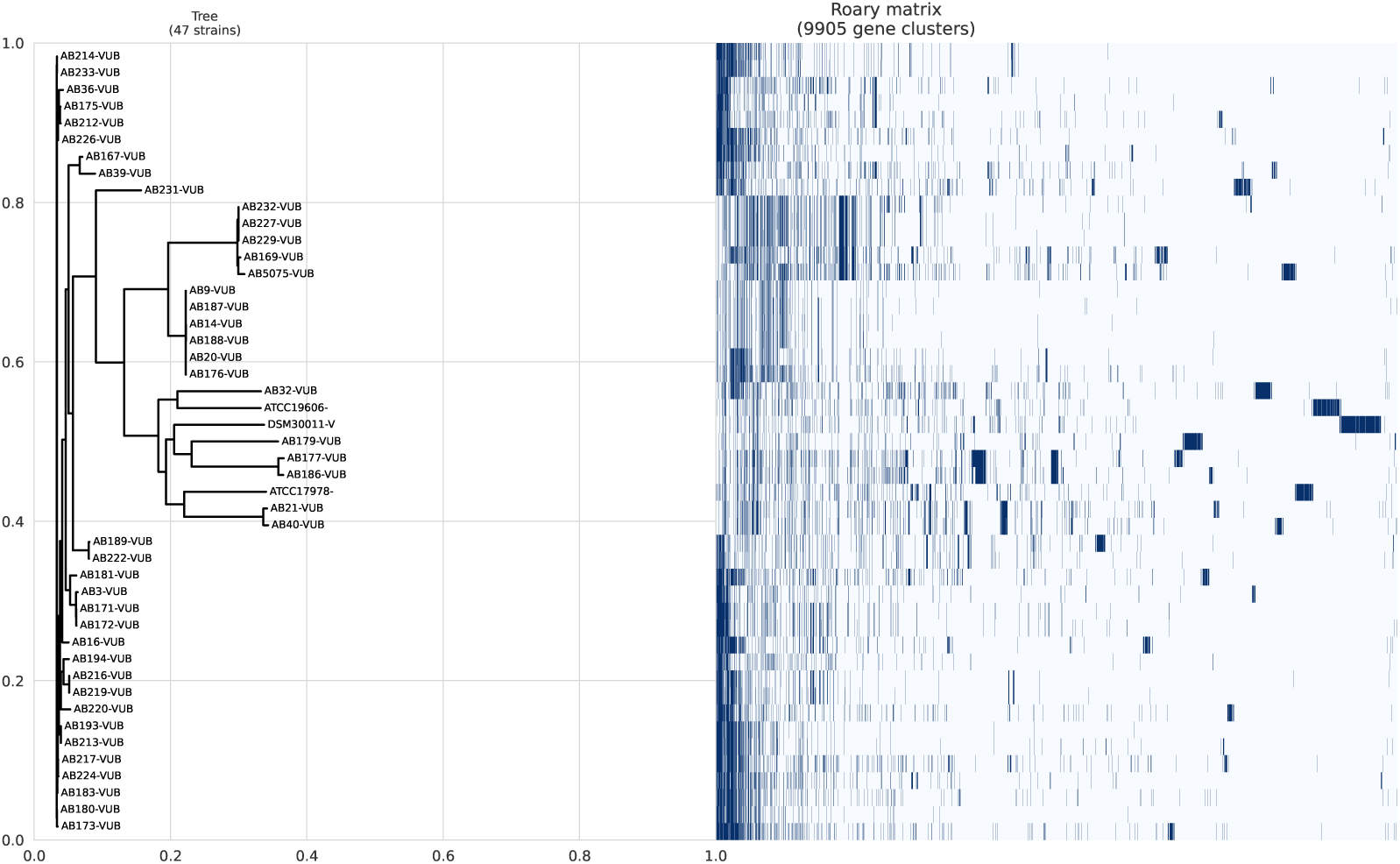
Pan-genome analysis of the complete genomic sequences of *A. baumannii*. The tree on the left was constructed based on accessory genome elements. The presence (in blue) and absence (in white) of accessory genome elements are displayed on the right.

**FIG 2.**
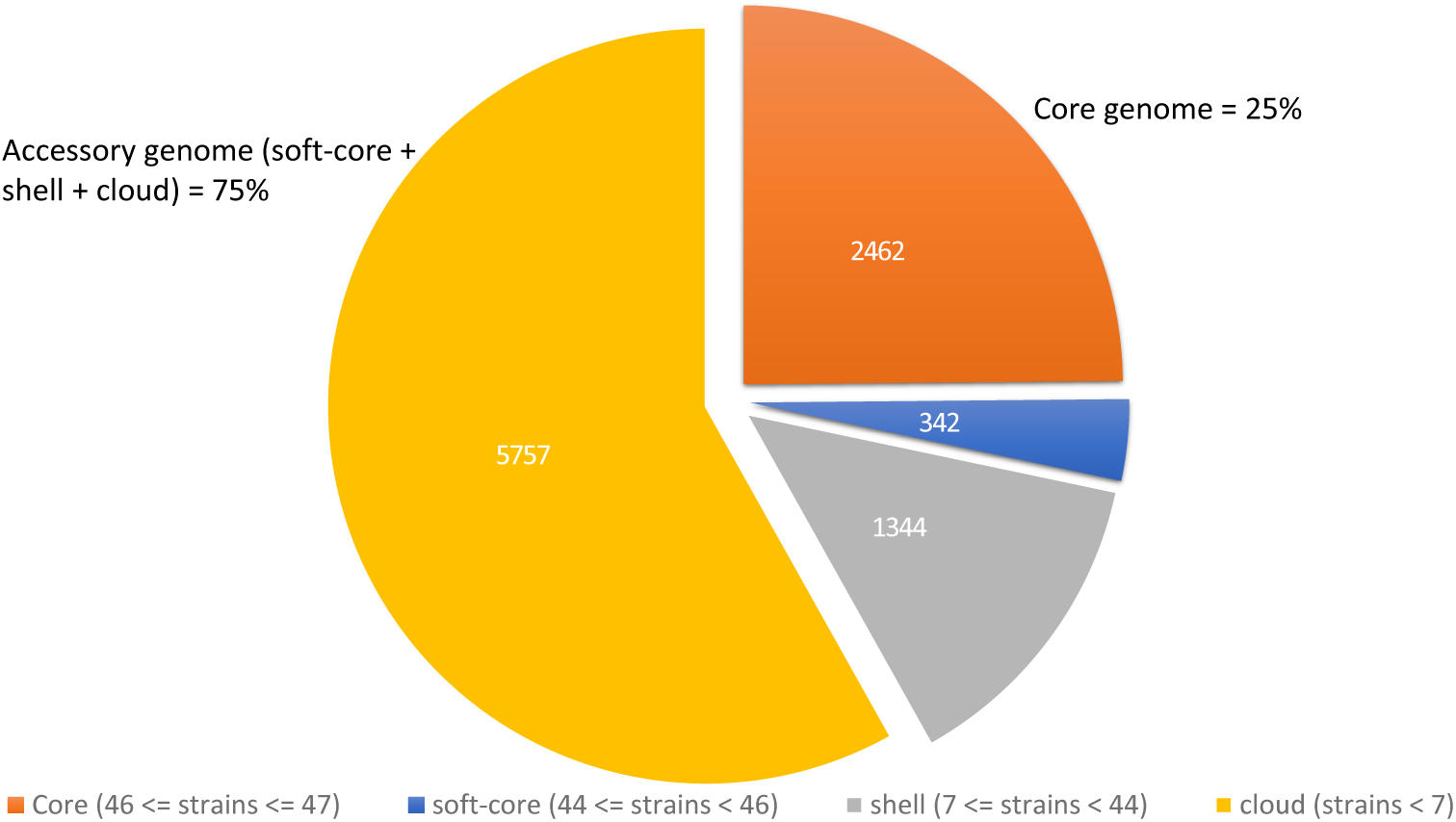
Pie chart of the pan-genome showing the distribution of core and accessory genes. Accessory genes are further divided into soft-core, shell, and cloud categories.

### 16S rRNA-based delimitation

Species delimitation based on 16S rRNA sequences produces different clustering patterns with ABGD and ASAP methods. ABGD grouped all sequences into a single cluster. In contrast, ASAP detected meaningful variation by identifying two groups in its best partition. All 16S rRNA gene copies from strain AB179-VUB were consistently assigned to Group 2. Strains such as AB186-VUB and AB39-VUB showed intra-strain heterogeneity, with only specific 16S rRNA copies (e.g., copy3 of AB186-VUB and copies 3 and 5 of AB39-VUB) grouped with AB179-VUB, while their remaining copies fell into Group 1.

### CG-based species delimitation

ABGD, when applied to the core-genome alignment, identified 11 distinct groups (Figure **3**, **10**). In contrast, ASAP detected up to 23 groups (Figure **4**, **10**).

**FIG 3.**
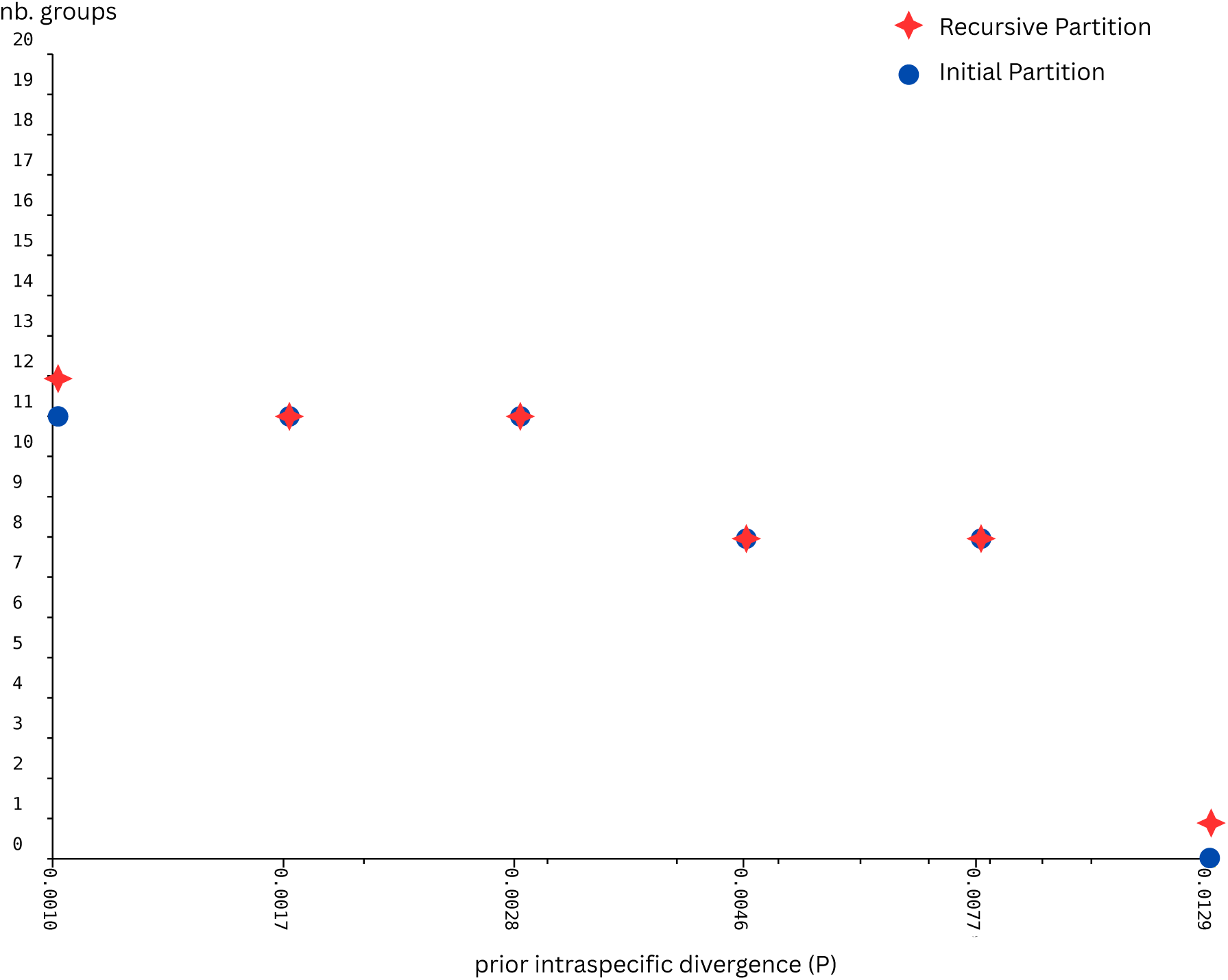
Species delimitation results based on core genome analysis using the ABGD method. Graph illustrating the number of groups identified as a function of prior intraspecific divergence (P), comparing initial and recursive partitions.

**FIG 4.**
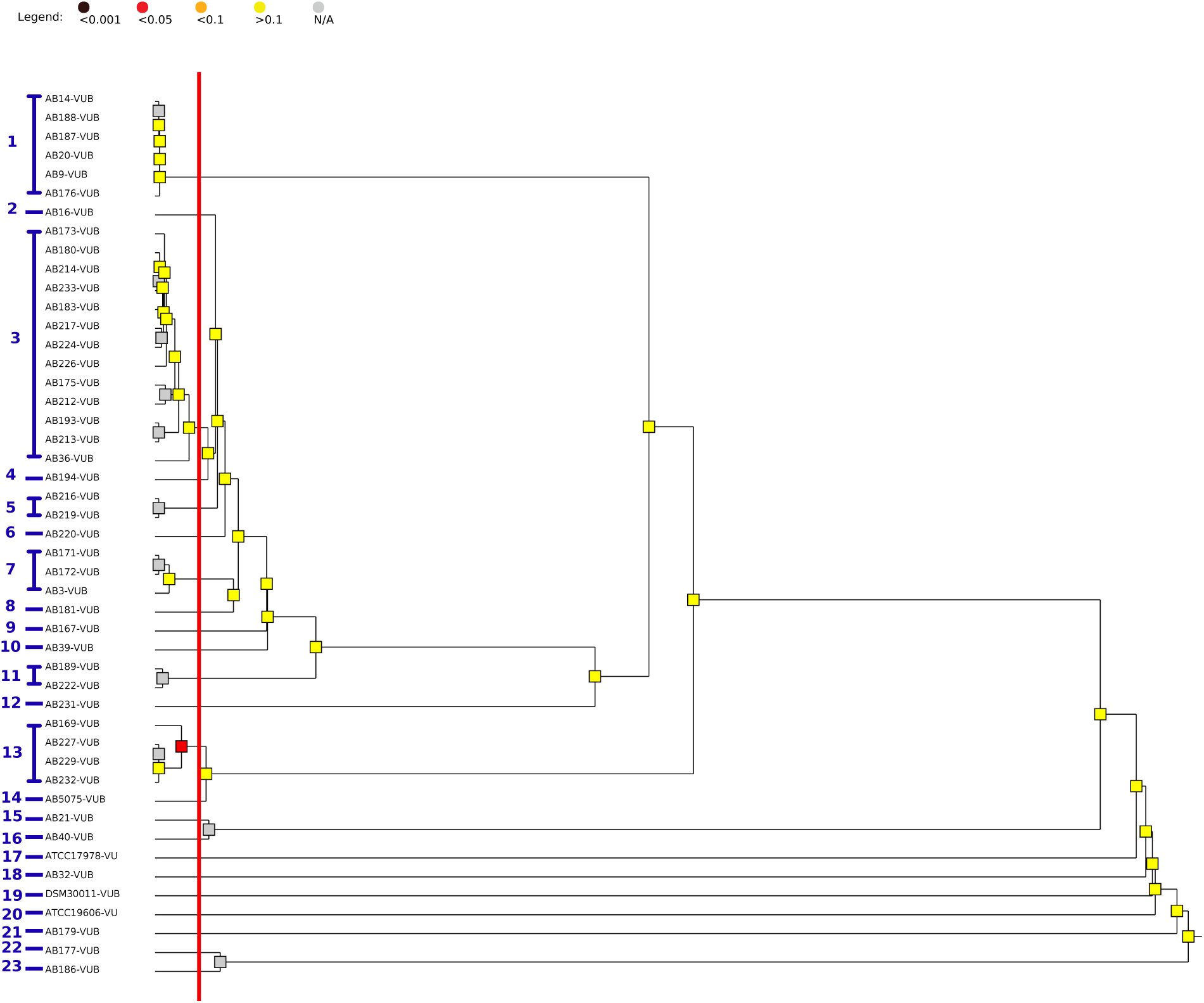
Species delimitation results based on core genome analysis using the ASAP method. Graphical representation of clusters identified by ASAP, with colored bars indicating the subsets.

**FIG 5.**
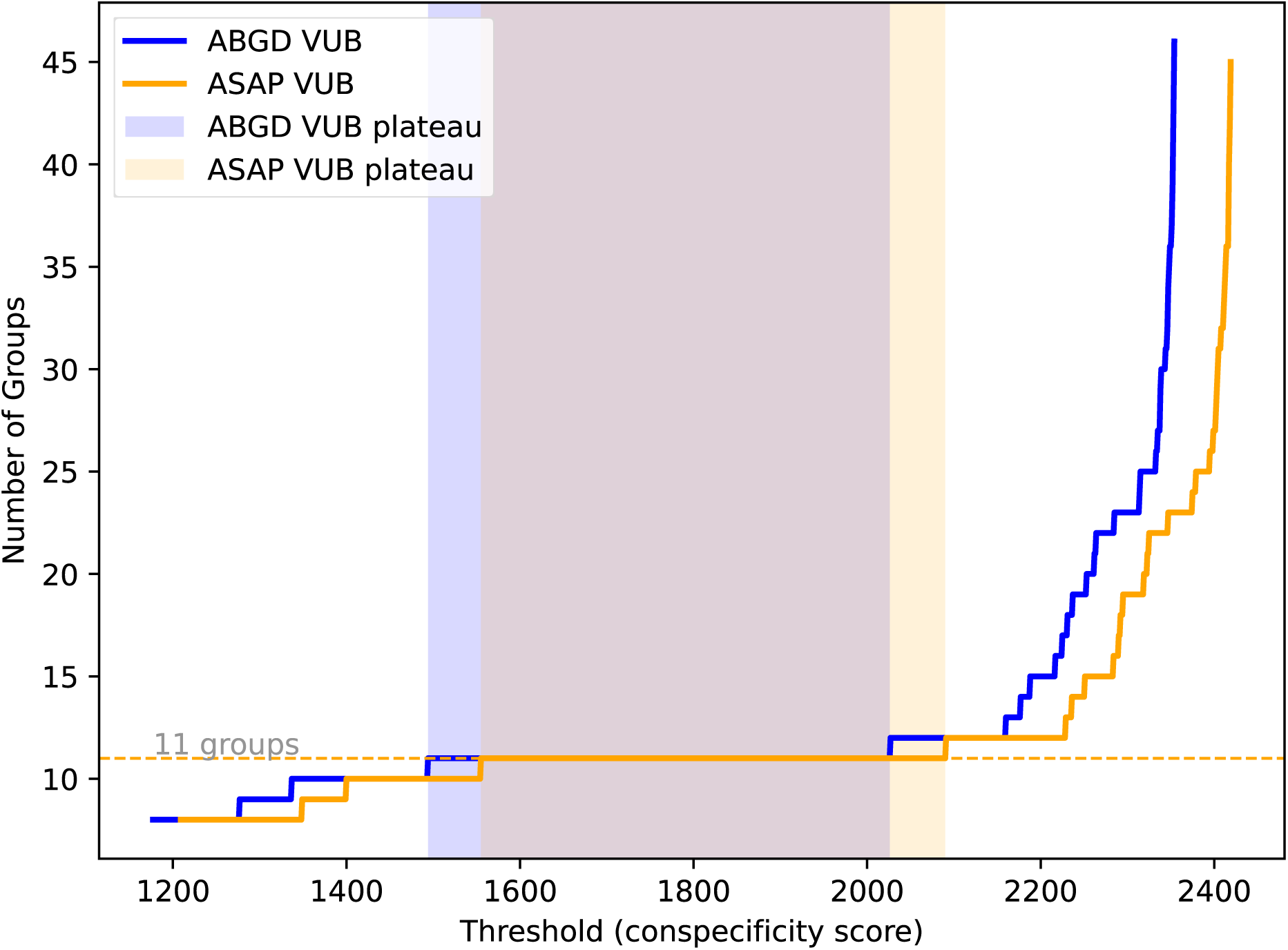
Number of CGCD-defined groups across co-occurrence thresholds in 47 VUB *A. baumannii* strains using ABGD and ASAP.

### ANI analysis

ANI was computed for all isolate pairs (Figure **6**). Although most ANI values exceeded 97.5%, applying a stricter 99.5% threshold delineated subgroups that aligned with the 11 clusters discovered by our ABGD CG analyses.

**FIG 6.**
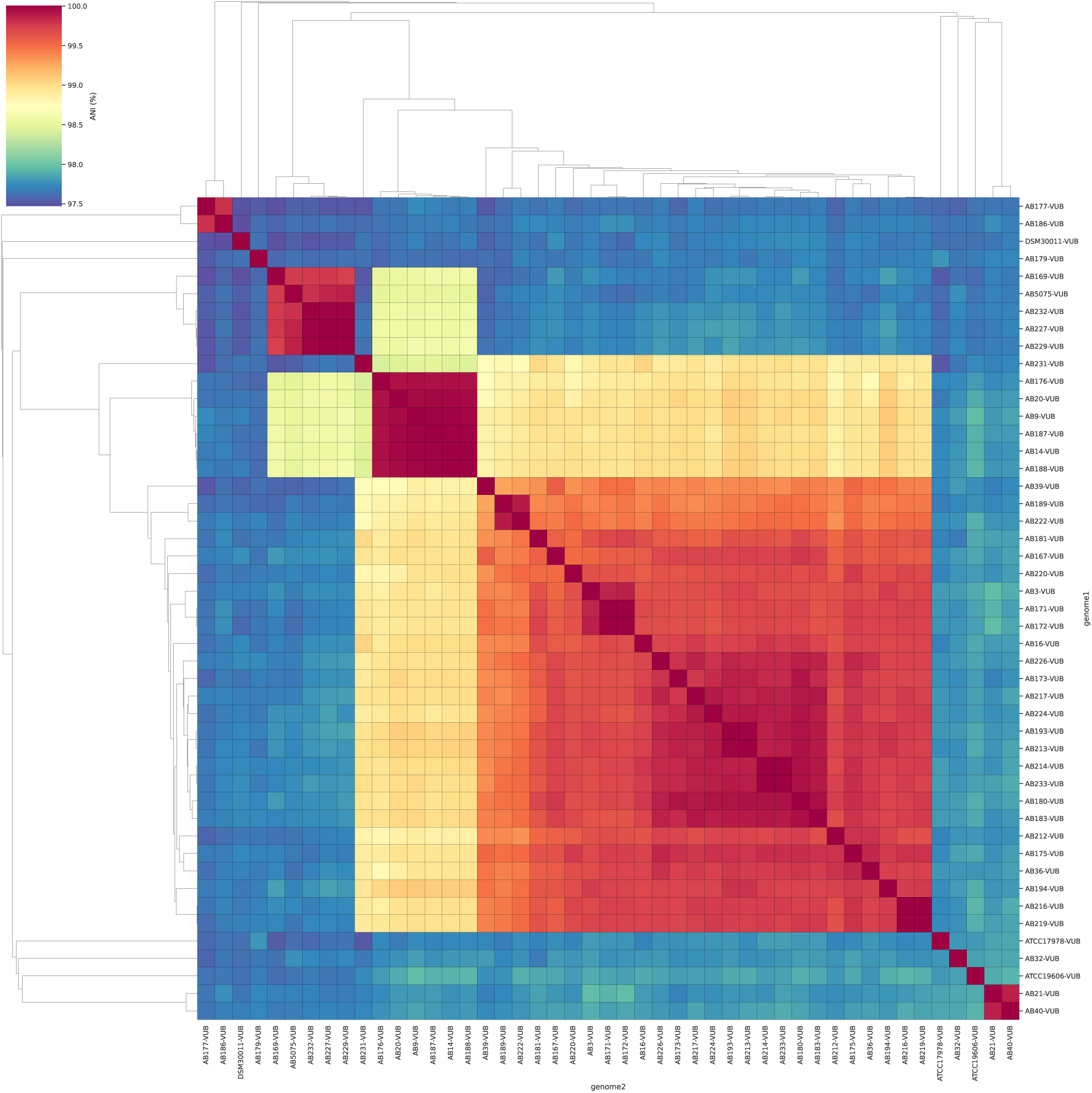
ANI analysis: Visualization of genomic similarities among the studied isolates.

### CGCD approach

The CGCD conspecificity matrices generated from ABGD (Figure **7**) and ASAP (Figure **8**) displayed well-defined block structures along the diagonal, each block corresponding to a distinct group of strains. Both methods produced a similar overall grouping pattern.

**FIG 7.**
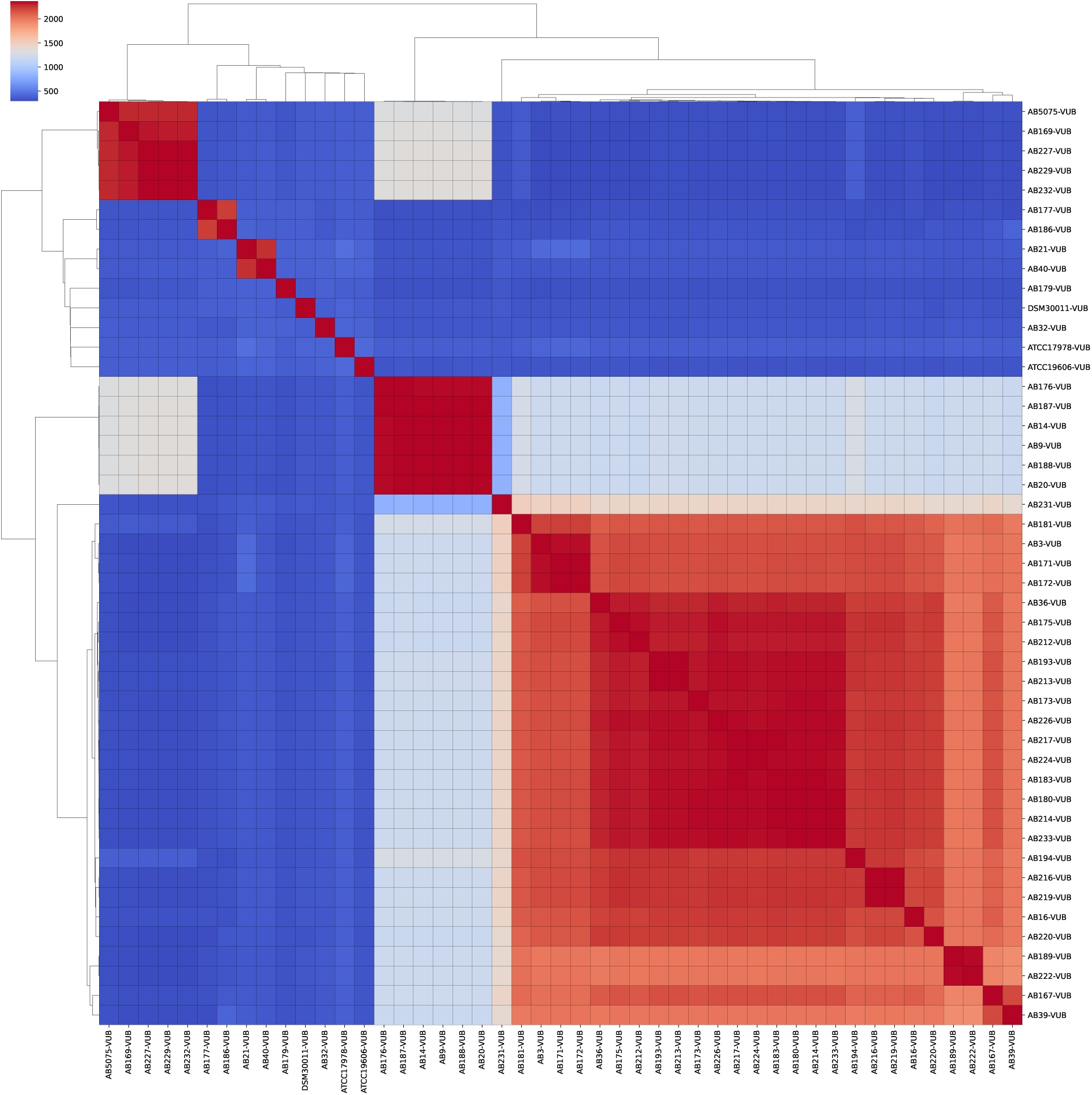
CGCD species delimitation results using the ABGD method.

**FIG 8.**
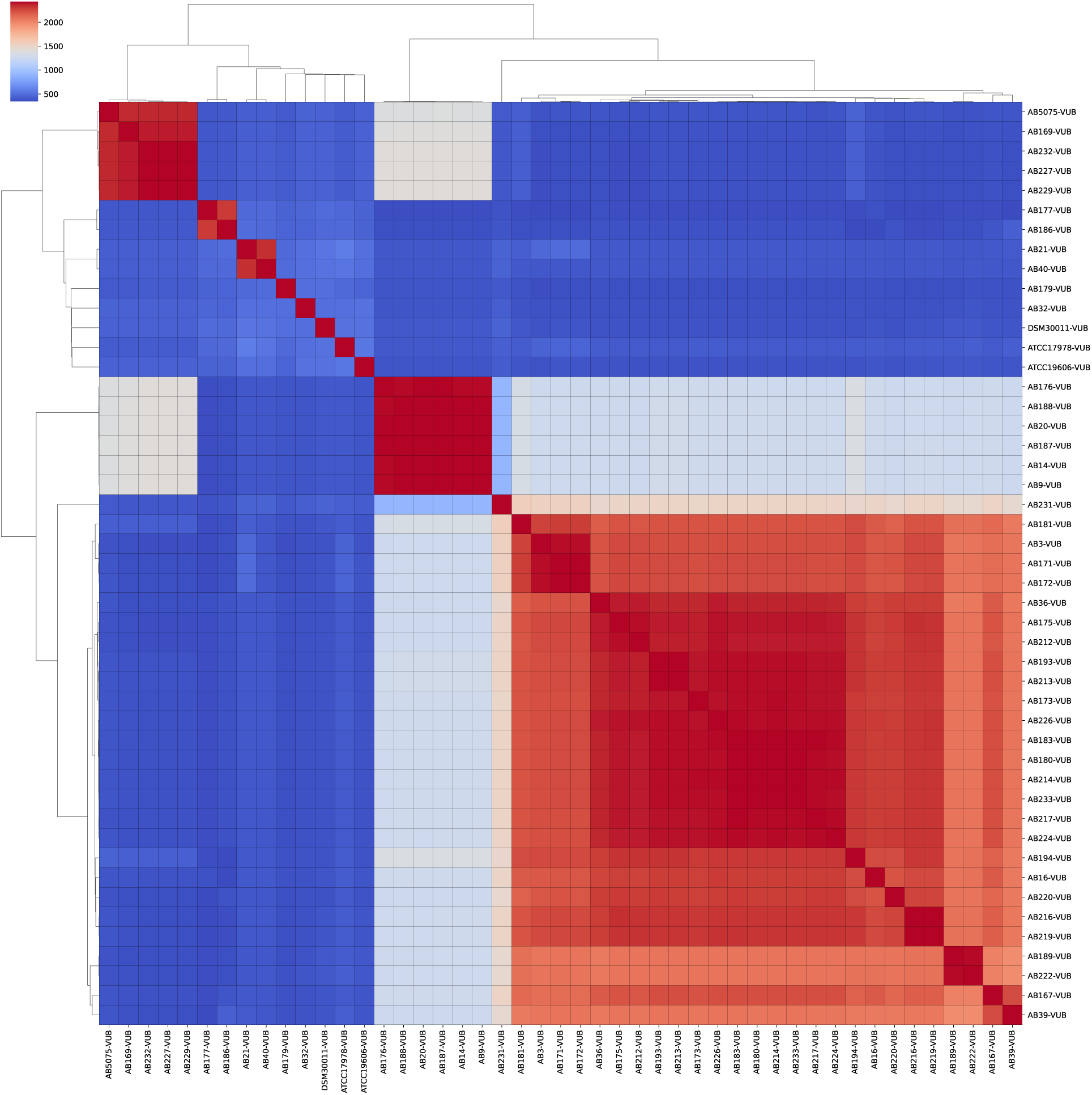
CGCD species delimitation results using the ASAP method.

When applying the threshold analysis, the curve (Figure **5**) plotting the number of groups against the co-occurrence threshold displayed a stability plateau. The stable value on this plateau corresponded to 11 groups, which was adopted as the final delimitation.

### Large-scale validation of genomic groups

To evaluate the robustness of genomic group delimitation inferred using the CGCD method, we examined the number of inferred groups containing the 47 VUB strains when embedded within the full dataset of 856 *A. baumannii* genomes across a range of co-occurence thresholds, defined as the number of core genes supporting conspecificity between genome pairs (Figure **9**). For both ABGD- and ASAP-based implementations of the CGCD framework, the number of inferred groups remained stable at 11 across a broad interval of threshold values. This stability is reflected by extended plateau regions in which group assignments were invariant despite changes in the conspecificity threshold. Outside these intervals, lower thresholds resulted in group merging (under-splitting), whereas higher thresholds led to progressive group fragmentation (over-splitting).

**FIG 9.**
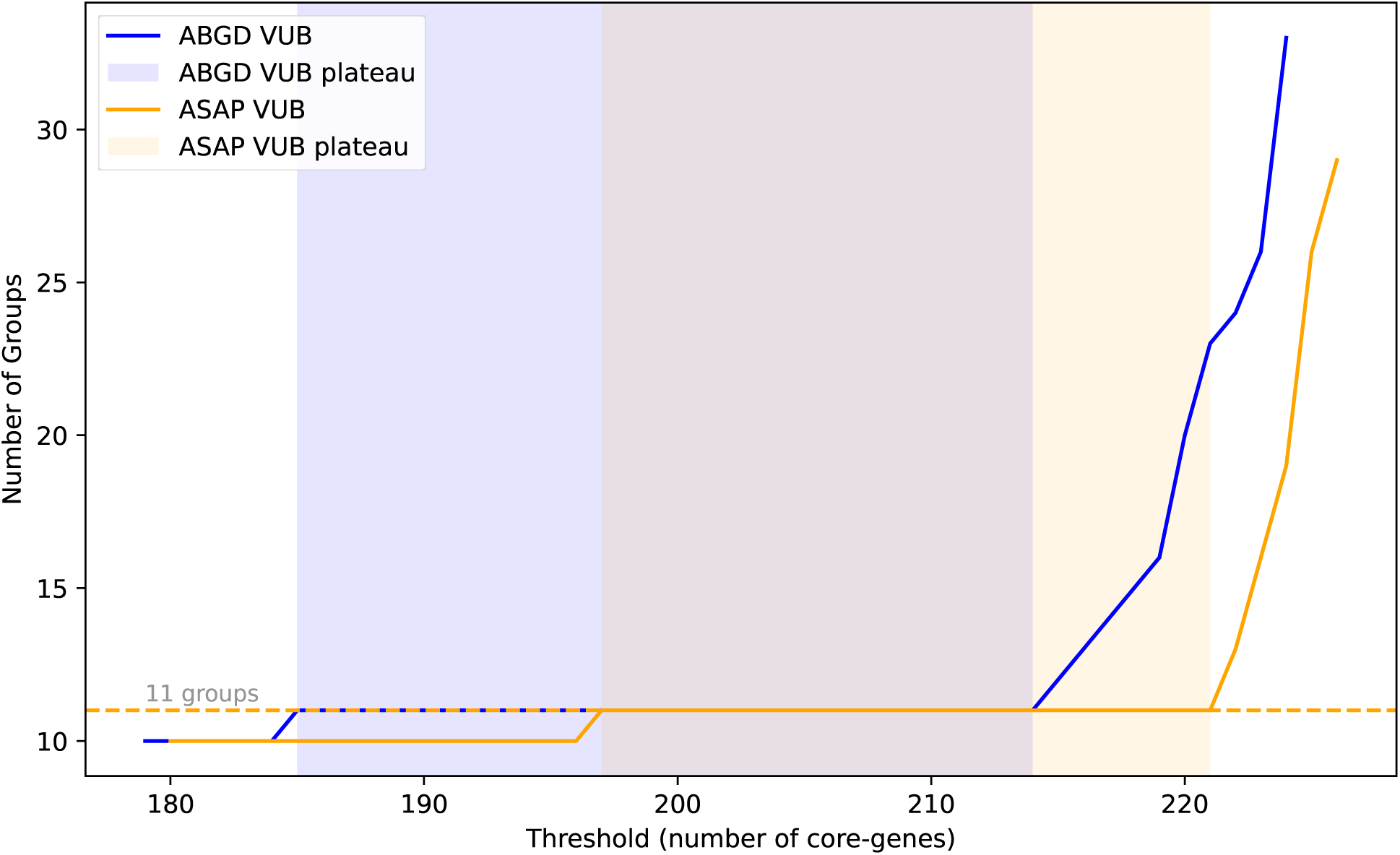
Validation of the best CGCD-defined groups from 47 VUB *A. baumannii* strains in a larger dataset of 856 genomes.

**FIG 10.**
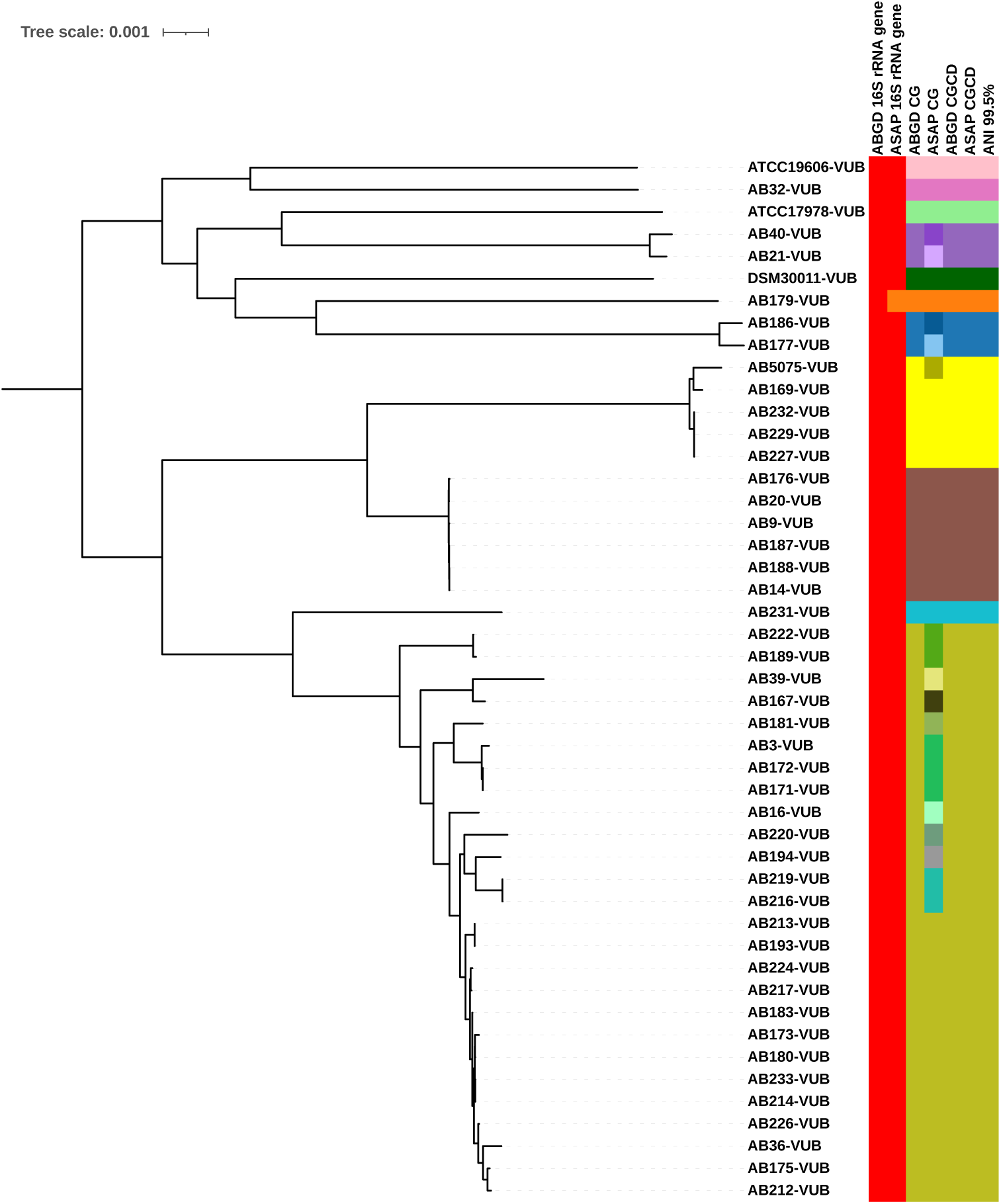
Phylogenetic tree of the core genome with group classification according to the applied analysis methods. CG: Core Genome, CGCD: Core-gene Consensus Delimitation.

### Phylogenetic and accessory genes analysis

We further evaluated the 11 CGCD groups using phylogenetic and accessory genome analyses. Mapping the groups onto a core-genome phylogeny revealed that each forms a distinct clade (Figure **10**), indicating that these clusters represent evolutionarily independent lineages rather than random groupings. We also constructed a clustering tree based on the presence–absence matrix of accessory genes (Figure **11**). In this tree, all strains grouped consistently with the CGCD assignments, with the exception of a single strain, AB231-VUB, which appeared within group 11.

**FIG 11.**
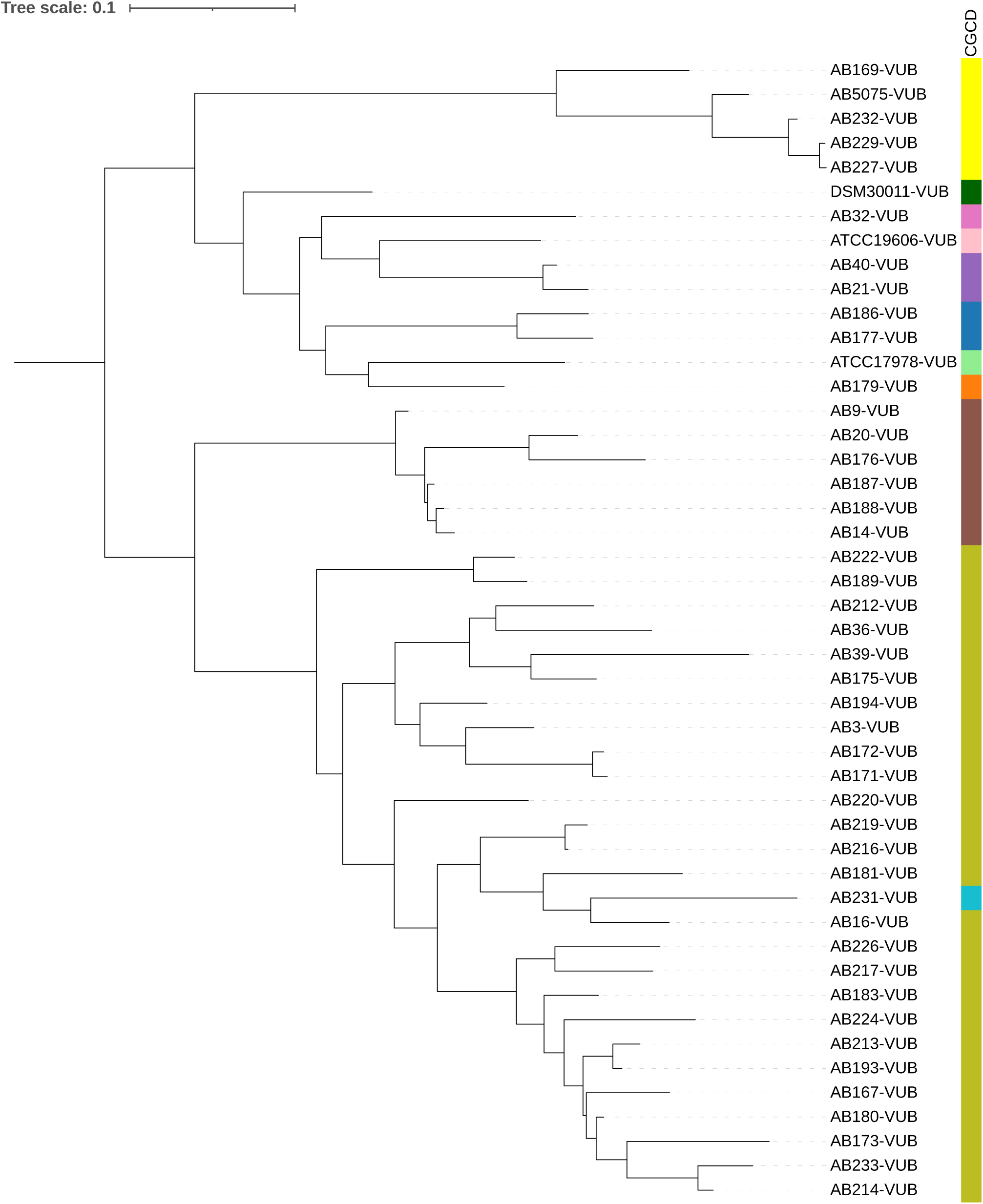
Clustering tree of the accessory genome with group classification according to the CGCD method. CGCD: Core-gene Consensus Delimitation.

## Discussion

Our findings highlight the extensive genomic diversity of *A. baumannii*, primarily driven by accessory genes associated with antibiotic resistance and virulence factors. Core genes account for only about 25% of the total genome, a proportion significantly lower than in other species, such as 59% in *Campylobacter* [34], 60% in *Neisseria* [35], and 88% in *Pseudomonas aeruginosa* [36]. This indicates that *A. baumannii* has a remarkable ability to acquire and discard genes through horizontal gene transfer [6, 37], facilitating adaptation to a wide range of environments, particularly clinical settings with intense antibiotic pressures.

Regarding species delimitation, our multifaceted approach demonstrated that methods based solely on 16S rRNA gene remain inadequate for resolving intraspecific diversity in *A. baumannii*. In our dataset, ABGD grouped all sequences into a single cluster, whereas ASAP identified only one distinct variant –AB179-VUB– assigning it to a separate group. This finding is consistent with the recognized low resolution of 16S rRNA for distinguishing closely related strains.

In contrast, applying ABGD and ASAP directly to the concatenated core-genome (CG) alignment yielded method-dependent results. ABGD identified 11 genomic groups, whereas ASAP produced a finer partition into 23 groups. However, concatenated CG analyses still treat all loci as a single evolutionary unit, potentially masking locus-specific divergence signals.

ANI analyses provided additional context for these patterns. Most pairwise ANI values exceeded 97.5%; however, applying a stricter cut-off of 99.5% still recovered the same 11 genomics groups identified by ABGD CG. This is consistent with pervious observations that ANI gaps between 99.2% and 99.8% can reflect meaningful intra-species structuring [25, 38]. Under conventional ANI-based criteria, the 11 groups identified here would thus be interpreted as subgroups rather than distinct species. This highlights an inherent limitation of ANI: by averaging nucleotide identity across the entire genome, ANI may lack the resolution to detect fine-scale population structure and can obscure locus-specific divergence signals. Alternatively, these groups may represent emerging evolutionary lineages that are in the early stages of divergence and have not yet crossed established ANI thresholds. In either scenario, ANI alone is insufficient to fully capture the underlying population structure, whereas gene-by-gene approaches such as CGCD provide complementary biological insight that may be overlooked by whole-genome similarity metrics.

The Core-gene Consensus Delimitation (CGCD) approach, extends ABGD and ASAP methods to a multi-locus framework by analyzing each core gene individually and integrates the resulting partitions into a conspecificity matrix. By scanning co-occurrence thresholds from 50% to 100% and visualizing the number of groups as a function of the threshold, we identified a clear stability plateau. The stable value within this plateau corresponded to 11 robust groups, which we adopted as the final delimitation. This threshold-guided step ensured that the chosen grouping was not an artifact of a single arbitrary cut-off, but instead reflected a consensus signal consistently supported by many independent loci. Importantly, the 11 CGCD groups corresponded closely with those detected by ABGD CG and ANI at 99.5%.

The robustness of this delimitation was further supported by large-scale validation using all publicly available *A. baumannii* whole genomes at the time of analysis. When the CGCD framework was applied to the expanded dataset of 856 genomes, the same 11 genomic groups were recovered, with group boundaries remaining stable despite the substantial increase in sampling breadth and genomic diversity. These clusters are differentiated across multiple levels: genomically, by CGCD analysis (Figure **5**); evolutionarily (Figure **10**), as each forms a distinct clade in the core-genome phylogeny; and functionally (Figure **11**), based on the presence-absence matrix of accessory genes. Together, these lines of evidence support the interpretation that the groups represent biologically meaningful, separate species.

Beyond the scope of *A. baumannii*, the CGCD framework is universally applicable, representing the first species delimitation method that can be applied across the tree of life, from bacteria to animals and plants. By capturing independent gene-by-gene signals and integrating them into a consensus-based analysis, CGCD provides a reproducible, data-driven approach for robust species delimitation.

## Conclusion

In conclusion, our analyses demonstrate that the 11 CGCD-defined genomic groups within *A. baumannii* are consistently supported across multiple levels of evidence. They are genomically distinct, evolutionarily independent, and functionally differentiated, reflecting robust, biologically meaningful population structure. The CGCD framework not only resolves fine-scale diversity that is obscured by traditional ANI or single-marker approaches but is also universally applicable across the tree of life, providing a reproducible, data-driven method for species delimitation. These findings highlight both the power of CGCD and the existence of perviously underappreciated structure within *A. baumannii*, with important implications for microbial systematics, evolutionary studies, and pathogen surveillance.

## ACKNOWLEDGMENTS

DATA AVAILABILITY STATEMENT FUNDING

KELM is funded by BLU-ULB (Brussels Laboratory of the Universe) JFF is funded by… AV is a recipient of a junior postdoctoral fellowship of the Research Foundation – Flanders (FWO; file number 1287223N).

## Supplemental material

